# Mechanisms of gene death in the Red Queen race revealed by the analysis of *de novo* microRNAs

**DOI:** 10.1101/349217

**Authors:** Guang-An Lu, Yixin Zhao, Ao Lan, Zhongqi Liufu, Haijun Wen, Tian Tang, Jin Xu, Chung-I Wu

**Affiliations:** State Key Laboratory of Biocontrol, School of Life Sciences, Sun Yat-sen University, Guangzhou 510275, Guangdong, China D; Center for Personal Dynamic Regulomes, Stanford University, Stanford, California, USA; Department of Ecology and Evolution, University of Chicago, Chicago, Illinois 60637, USA

**Keywords:** *de novo* gene, new gene, microRNA, gene death, Red-Queen effect

## Abstract

The prevalence of *de novo* coding genes is controversial due to the length and coding constraints. Non-coding genes, especially small ones, are freer to evolve *de novo* by comparison. The best examples are microRNAs (miRNAs), a large class of regulatory molecules ~22 nt in length. Here, we study 6 *de novo* miRNAs in *Drosophila* which, like most new genes, are testis-specific. We ask how and why *de novo* genes die because gene death must be sufficiently frequent to balance the many new births. By knocking out each miRNA gene, we could analyze their contributions to each of the 9 components of male fitness (sperm production, length, competitiveness etc.). To our surprise, the knockout mutants often perform better in some components, and slightly worse in others, than the wildtype. When two of the younger miRNAs are assayed in long-term laboratory populations, their total fitness contributions are found to be essentially zero. These results collectively suggest that adaptive *de novo* genes die regularly, not due to the loss of functionality, but due to the canceling-out of positive and negative fitness effects, which may be characterized as “quasi-neutrality”. Since *de novo* genes often emerge adaptively and become lost later, they reveal ongoing period-specific adaptations, reminiscent of the “Red-Queen” metaphor for long term evolution.

## Introduction

Haldane (1932) may be the first person to suggest that new genetic materials might be continually generated. In the modern literature, genes can be of either of two different origins. The first class is derived from existing genic sequences by gene duplication, retroposition, elongation, fusion etc. Since its proposal (Haldane 1932), the hypothesis has been formalized and pursued by Susumu Ohno (Ohno 1970) and others (Li 1982; Lynch and Conery 2003; Zhang 2013). The second class is new genes that originated from non-genic sequences, referred to as “*de novo*” genes (Chen et al. 2013). *De novo* genes that code for new proteins have been considered rare, and even improbable (Ohno 1970). Francois Jacob suggested that ‘the probability that a functional protein would appear *de novo* .. is practically zero’(Jacob 1977). Given the constraints on proteins’ length and structure, such reservations are at least understandable.

In the last decade, evidence on *de novo* protein-coding genes has been presented (Tautz and Domazetlošo 2011; Zhao et al. 2014; Schlötterer 2015) but controversies remain unresolved (Moyers and Zhang 2015; Mclysaght and Hurst 2016; Moyers and Zhang 2016; Domazetlošo et al. 2017; Moyers and Zhang 2017). As the skepticisms toward *de novo* genes are mainly about forming functional peptides, *de novo* noncoding genes may be more plausible without the demand for coding. The constraints could be further relaxed if the noncoding gene is short.

In this backdrop, *de novo* microRNAs (miRNAs) that function in post-transcriptional repression of target mRNAs(Bartel 2004) would be particularly relevant. First, mature miRNAs (miRs) are short, only 22 nt long, and the biogenesis requires only a loosely formed hairpin structure of < 100 nt. It has been estimated that a *Drosophila* genome can easily form more than 10,000 hairpins that could potentially yield mature miRs (Lu et al. 2008). Second, unlike putative coding genes whose functions are difficult to assay, functional *de novo* miRNAs should be able to repress their target mRNAs. Third, many *de novo* miRNAs have been proposed and verified (Meunier et al. 2013; Lyu et al. 2014; Mohammed et al. 2014). Therefore, miRNAs could be considered a paradigm for the evolution of *de novo* genes.

The evolutionary dynamics of *de novo* genes, coding or noncoding, is of great interest. Since much of the genome of higher eukaryotes has no known function, the occurrence of functional genes in these regions would have significant impacts on genome organization. Nevertheless, *de novo* genes, if common, present a dilemma - the constant influx of *de novo* genes would mean steady turnover of the genome, which should be evident between species of the same taxa, say, among mammals. Since the rapid turnover does not appear true, a reasonable conjecture would be that *de novo* genes often emerge adaptively but die subsequently(Lyu et al. 2014).

We select for detailed analysis a subset of *de novo* miRNAs in *Drosophila* that have been shown to evolve adaptively after emergence (Lyu et al. 2014). Their subsequent evolution would be a test of the Red Queen hypothesis, which posits adaptation as a never-ending process (Van Valen 1973; Salathé et al. 2008; Liow et al. 2011). In this view, an adaptive *de novo* gene might quickly lose out if it cannot keep up with all other evolutionary changes in an unfamiliar landscape. Whether, how and why new genes die is the central question of this study.

## Results

### Evolution of *de novo* miRNA genes in *DrosoPhila*

The hallmark of *de novo* genes is their presence in a well-defined clade of closely related species and absence outside of it. Furthermore, the time frame relative to the level of divergence has to be short such that the absence is not confounded by “the failure to find”, a common source of controversy (Moyers and Zhang 2015; Moyers and Zhang 2016). A densely populated phylogenetic tree of species with relatively small genomes is best for new gene analysis. The phylogeny of *Drosophila* is thus ideal for studying *de novo* miRNAs (Fig. 1A).

A more challenging issue is the function of the putative *de novo* genes (Tautz and Domazetlošo 2011; Wu and Zhang 2013; Mclysaght and Hurst 2016). Many *de novo* miRNAs reported may have never been functional (Lu et al. 2008), or to use a term borrowed from the study of retroposed genes, “DOA (dead-on-arrival;) (Petrov et al. 1996; Petrov and Hartl 1998)”. Different criteria have been employed to exclude “DOA” miRNAs (Lu et al. 2010).

In a previous study of ours (Lyu et al. 2014), *de novo* genes are restricted to miRNAs that are not neutrally evolving in their DNA sequences. They reported many younger miRNAs (< 30 Myrs old) that show the adaptive signature in sequence evolution. By such criteria, *de novo* miRNAs are generally X-liked and testis-specific in expression (see the supplemental text S1 for details), which are also the characteristics of newly formed coding genes. Furthermore, more than half of *de novo* miRNAs are found in clusters. The *mir-982* and *mir-972* clusters, both X-linked as shown in Fig.1B, stand out. The *mir-982* cluster contains only members <30Myrs in age, while *mir-972s* have older ones (Fig. 1B). Here, we choose 6 of the more highly expressed (Supplemental Table S1) and rapidly evolving miRNAs from three age groups for detailed analysis (different colors in Fig. 1B denoting the age of < 4 Myrs, 4 - 30 Myrs and 60-250 Myrs).

### Characterization of miRNA knock-out lines: Sequencing and transcriptome assays

Using the TALEN technology (Boch 2011), we generate knockout lines for the 6 miRNAs shown in Fig. 1A (see Methods, Supplemental Table S2;Supplemental Fig.S1). Among them, *mir-973*, *mir-975* and *mir-977* are older, emerging 60 - 250 Mrys before present. The remaining three are *mir-978*, *mir-983* (4 - 30 Mrys old) and *mir-984* (< 4 Myrs old). Each deletion has been confirmed by PCR and DNA sequencing (upper panels of Fig.1C-1H). Loss of the mature miRNA product is validated by qPCR on RNA extracts from the testis (Bottom-panels of Fig.1C-1H). We use this collection of miRNA deletions for the rest of the study.

**Figure 1.**
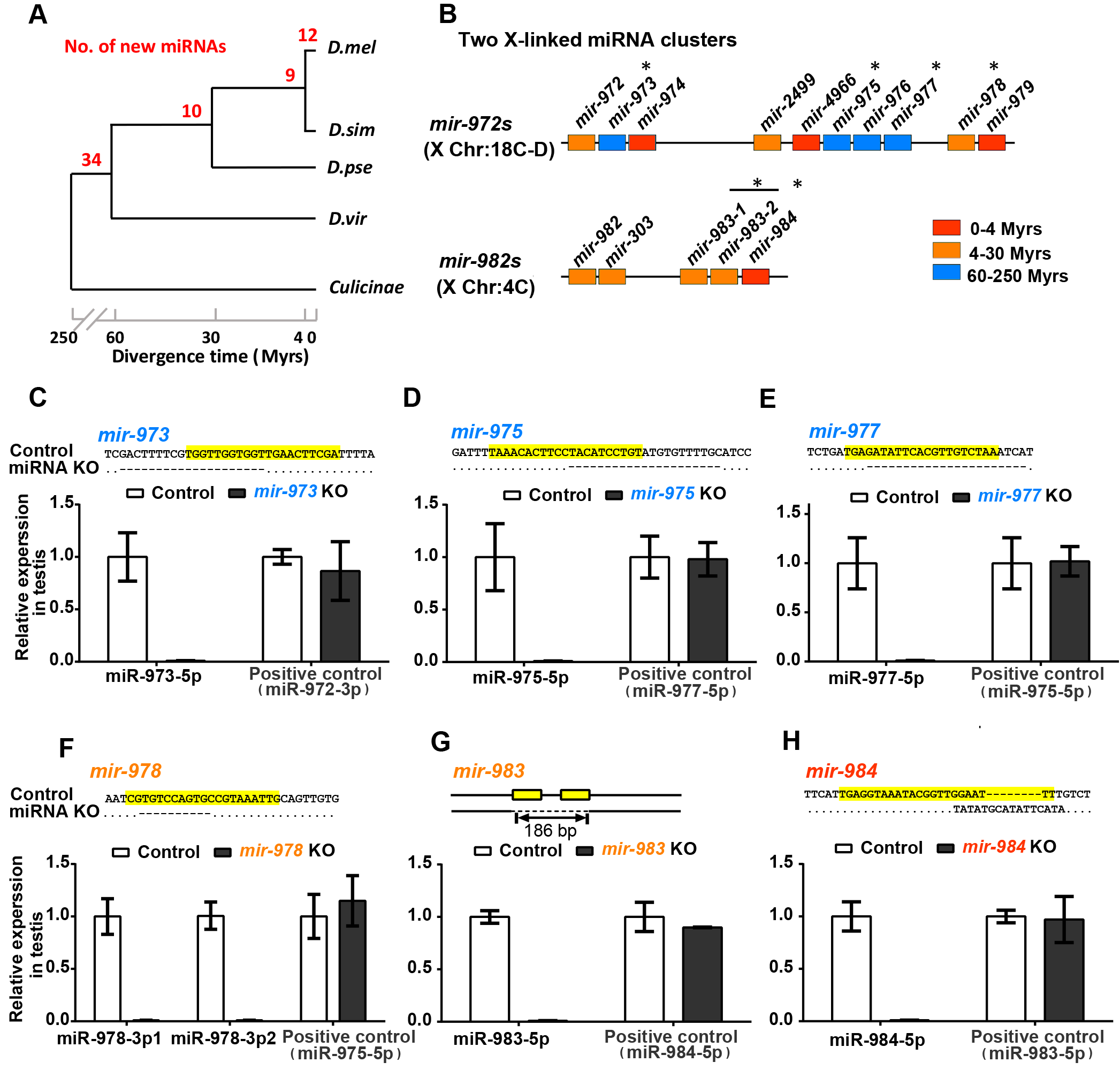
Emergence of *de novo* miRNAs and the creation of knockout (KO) lines. **(A)** The emergence of new miRNAs in *Drosophila*. The numbers in red indicate miRNA emergence in each time interval of *Drosophila* phylogeny. Note the “pyramid” age structure with many very young miRNAs suggesting many deaths before aging. The age distribution is sourced from our previous study(Lyu et al. 2014). *D. mel*: *D. melanogaster; D. sim: D. simulans; D. pse*: *D. pseudoobscura; D. vir*: *D. virilis*. **(B)** miRNA mutants in this study fall into two *de novo* miRNA clusters. *De novo* miRNAs tend to be in clusters and are X-linked (Lyu et al. 2014). The two clusters of our focus are shown here with * indicating the 6 knockout creations. Colors denote miRNAs’ age as indicated. **(C)** To **(H)** Verification of miRNA knock out efficiency. Upper panels: DNA sequences of miRNA KO strains with the mature sequences highlighted in yellow. Lower panels: relative expression of miRNAs in the testis. No miRNAs are detectable in the KO lines whereas the expressions of the neighboring miRNAs are normal (positive controls). Note that *mir-978* has two 5’iso-miRNAs (miR-978-3p1 and miR-978-3p2), which are processed from one precursor in the *D. melanogaster* testis.

The “molecular phenotypes” of a *de novo* miRNA can be assayed by its immediate impact on the transcriptome via the direct targets, an advantage of studying miRNAs over putative coding genes (Zhao et al. 2014; Gubala et al. 2017). RNA-seq measurements are done on the testes of flies bearing the miRNA deletion as well as those of the wild type flies. The quality of the transcriptome data is high as shown in Methods (Pearson’s r > 0.98 between replicates). Fig. 2A-2C presents the repression effects of the three younger miRNAs. Two (*mir-978* and *mir-983*) indeed show significant repressions on their predicted targets (Fig.2A-2B; *P* < 0.05 by the Kolmogorov-Smirnov test). The youngest miRNA (*mir-984*) that is less than 4 Myrs old does not show significant repression effects (Fig. 2C). Given the very young age, target prediction might be much less reliable than that for the older miRNAs.

Below, we analyze all repressions, direct or indirect, without requiring target predictions. Given their testis-specific expressions, one would expect the indirect targets to be enriched for genes previously characterized as male-biased (Gnad and Parsch 2006). The prediction is supported by the result shown in Fig. 2D. While the ratio of male-biased/non-based genes is 0.57, this ratio is > 1 for all target genes (direct and indirect) of each *de novo* miRNA (*P* < 0.01 in all cases). Thus, these *de novo* miRNA genes all have an impact on a specific subset of genes in the testis transcriptome

**Figure 2.**
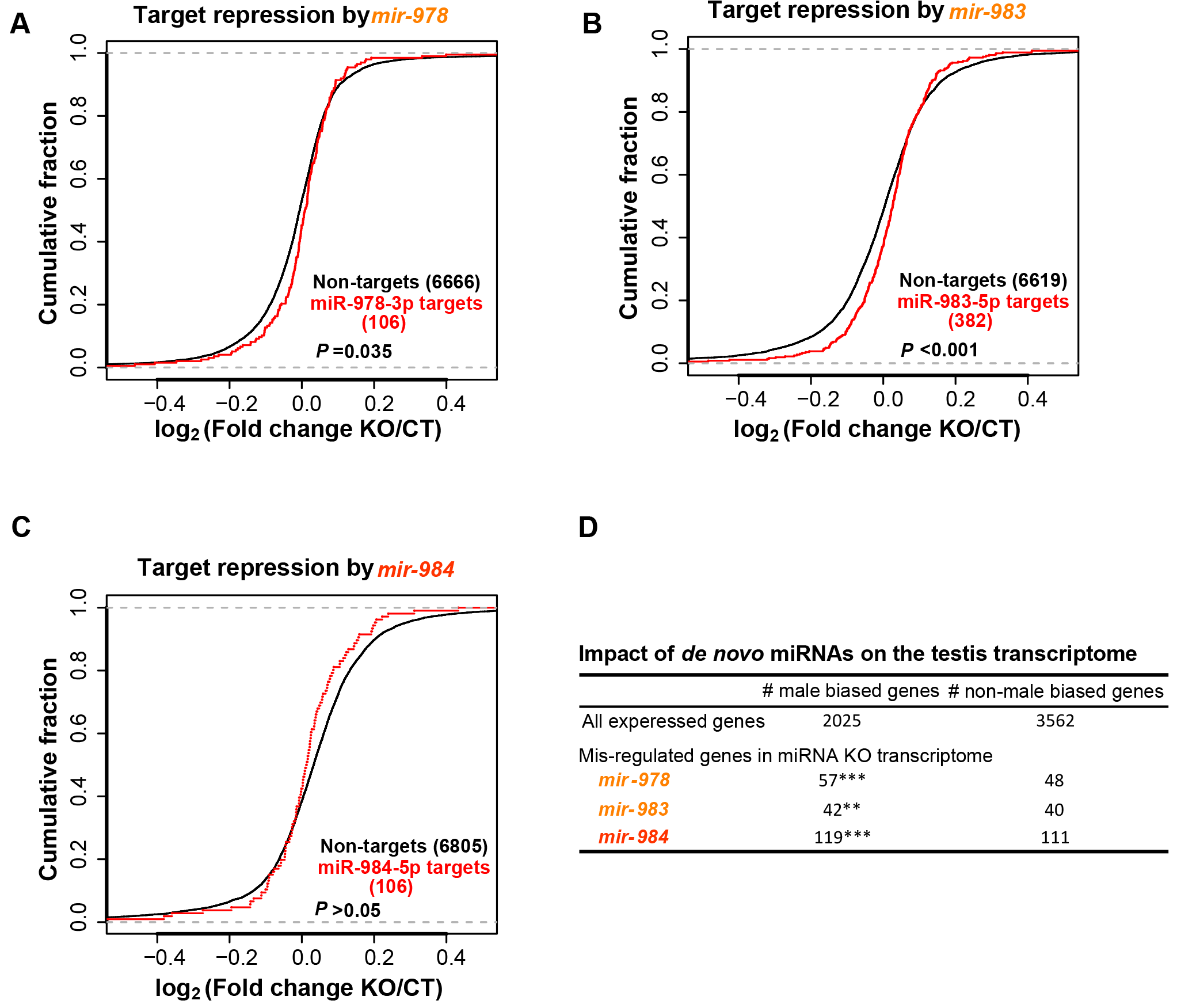
Impact of *de novo* miRNAs on target gene expression. **(A)** *mir-978* target repression. Target pools of the two *mir-978* iso-miRs overlap. Targets in this overlapped pool are significantly up-regulated in the *mir-978* KO line (P=0.035, Kolmogorov-Smirnov test). **(B)** *mir-983* target repression. Targets of *mir-983* (miR-983-5p) are significantly up-regulated compared to non-targets in the *mir-983* KO line (*P*<0.001, Kolmogorov-Smirnov test).**(C)** *mir-984* target repression. *mir-984* (miR-984-5p), the youngest of the miRNAs, do not significantly down-regulate their targets (*P* > 0.05, Kolmogorov-Smirnov test). **(D)** Impact of *de novo* miRNAs on the testis transcriptome. Genes mis-regulated in miRNA KO lines are enriched for male biased genes (Chi-square test, *P*<0.01 and *P*<0.001 marked ** and ***, respectively).

### Effects of *de novo* miRNAs on male fitness components

Given the testis-specific expression, we expect any fitness effect would be associated with male reproduction. Because miRNAs may concurrently affect multiple sub-phenotypes (Liufu et al. 2017). We assay the sub-phenotypes of male reproduction in the following order: 1) male fertility (progeny production), 2) sperm length and sperm competition, 3) males’ ability to repress female remating, 4) meiotic drive, 5) male viability, and 6) mating success of males. Although miRNAs (> 30 Myrs) have been expected to exhibit stronger fitness effects, they are found only marginally more so than the younger ones.

#### Male fertility (progeny production)

Progeny production is counted in two batches-in days 1-2 and in days 3-7 after mating (Supplemental Fig.S2). Each batch, accounting for about the same number of progeny, may reflect different attributes of male fecundity such as sperm transfer and sperm longevity in storage. In Fig. 3A and 3B, the absence of each of these miRNAs does not appear to have any significant effect on male fertility. In contrast, the three older miRNAs (*mir-973*, *mir-975* and *mir-977*) seem to be required in the later stage of progeny production. The three knockout lines have somewhat better performance than the control in the first two days but become worse in the subsequent 5 days. For the three younger miRNAs (*mir-978*, *mir-983*, *mir-984*), the differences between the two stages are less pronounced.

The detailed daily data for one older miRNA (*mir-977*) and a younger one (*mir-978*), which are adjacent in the same cluster, are shown in Fig. 3C-3F (For the rest, see Supplemental Fig.S3). For *mir-977* knockout males, the number of eggs and the number of progeny both show a decline in the Day 3 - 7 data (Fig.3D, Multiple t-tests with the Holm-Sidak correction, alpha=5%). For *mir-978*, Fig. 3E and **3F** show a trend in the opposite direction but the difference between the knockout and the control is not significant. Apparently, the older *mir-977* is needed for some aspects of sperm function whereas the younger *mir-978* appears dispensable for (or even harmful to) those same aspects.

**Figure 3.**
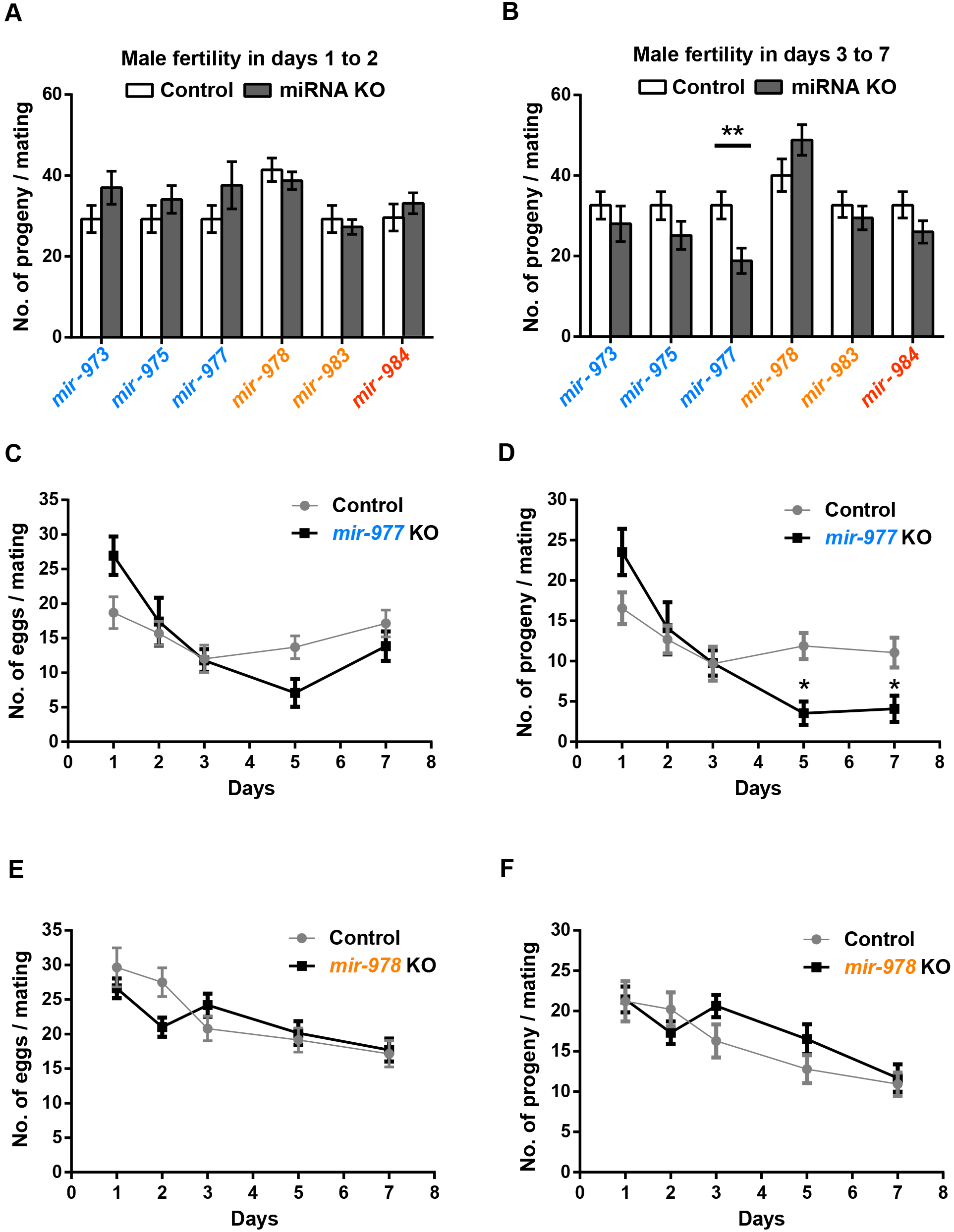
Relative fertility in miRNA KO lines. After mating with either wildtype or miR KO males, females are permitted to lay eggs for 7 days. **(A)** Number of progeny in the first two days. **(B)** Number of progeny in days 3-7. Overall, the fertility differences between wildtype and KO males are small. The three older miRs seem to have a larger impact on days 3-7 while the younger miRs do not show a consistent pattern. Except for *mir-977* (Student’s *t*-test: **, *P*<0.01), none of the differences is significant. **(C-D)** Day-to-day fertility analyses of *mir-977* for egg laying and progeny count, respectively. Mean±SEM is shown. Fertility differences between wildtype and KO males are significant in later days (Multiple t-tests with the Holm-Sidak correction, *P*=0.0009 on day five and *P*= 0.011 on day seven). **(E-F)** Day-to-day fertility analyses of *mir-978*. There is a slight but insignificant decrease during the first two days and a slight increase in subsequent days for the KO line.

#### Sperm length in relation to sperm competition

Progeny count is a measure of male fertility when a single male inseminates virgin females. It reflects sperm count but does not reflect sperm competitiveness. However, since wild-caught *Drosophila* females usually carry sperm from several males, sperm competitiveness may be no less an important measure of male fitness than the simple progeny count (Parker 1970; Griffiths et al. 1982; Jones and Clark 2003). An important indicator of sperm competitiveness is sperm length which has been shown to be positively correlated with the competitive edge (Miller and Pitnick 2002; Pattarini et al. 2006; Lüpold et al. 2016). Fig. 4A shows that, with the exception of the *mir-977* mutant, all other miRNA deletions result in longer sperm. In four of them, the length difference is statistically significant (Fig.4A, *P* < 0.05, Mann-Whitney *U* test). It is curious that the absence of these miRNAs appears to increase the sperm length.

The sperm of males with the miRNA knockout are not only longer but also more uniform in length. The coefficient of variation (standard deviation/mean) is consistently smaller than that in the wild type (Fig.4B). We suspect that both the mean and the variation in sperm length may contribute to sperm competition as reported below. It is curious that well established miRNAs often contribute to the canalization in development (Wu et al. 2009; Cassidy et al. 2013; Posadas and Carthew 2014). One would naturally have predicted that miRNA knockout would increase the cell-to-cell variation. The observations to the contrary may be another indication of the peculiarity of *de novo* miRNAs.

We now present the measurements of sperm competition, which has an offense and defense component. In the offense measure, females are mated to the control male and, after a resting period, are mated to a second male. This second male can be either the wild type male (control) or a male from the miRNA knockout line (experimental male). The proportion of the sperm from the second male is the offense measure. In the defense measure, the sequence of the first and second male is reversed and the progeny sired by the first male is the defense measure (See Methods, Supplemental Fig.S4). A higher score in either offense or defense indicates superior competitiveness.

Compared with the control, males carrying the knockout mutation in each of the three younger miRNAs have higher scores in both the offense and defense assays (Fig.4C-4D, *P* < 0.05, Mann-Whitney *U* test). Again, the absence of these genes unexpectedly improves the fitness. For the three older miRNAs the pattern is less consistent but the expectation of reduced fitness in knockout lines is not fulfilled. In Fig. 4E-4F, we observe the positive correlation between sperm competitiveness and sperm length. The correlation for the defense score is significant (*P* = 0.04). Since the offense score depends on many other parameters of male ejaculates (such as the content of the seminal fluid), the lower correlation between the offense score and sperm length is expected. Collectively, our data reveal the fitness improvement upon the deletion of the younger miRNAs. One would then expect their elimination by natural selection if other fitness components are equal. Obviously, other components are not equal as shown below.

**Figure 4.**
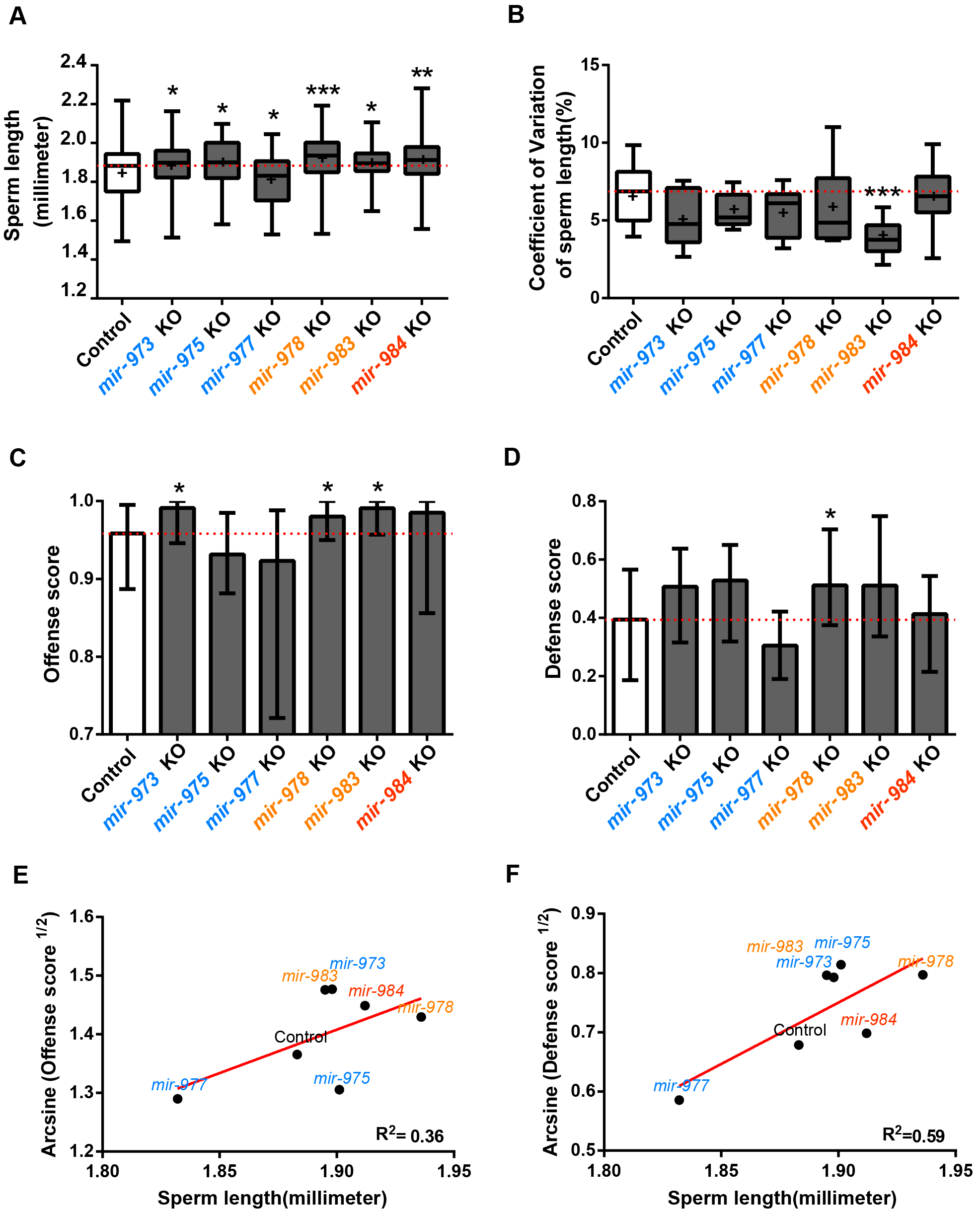
Sperm length and sperm competition. **(A)** Sperm length of miRNA KO and control males. Sperm length is recorded for 9 - 11 individuals with 9 - 13 sperm measured per individual. Except *mir-977*, all miRNA KO lines have significantly longer sperm compared to the control (Mann-Whitney *U* test: *, *P*<0.05, **, *P*<0.01, ***, *P*<0.001).**(B)** Coefficient of variation (standard deviation / mean) of sperm length. All miRNA KO males tend to have sperm of more homogeneous length, especially *mir-983* KO (Mann-Whitney *U* test, ***, *P*<0.001).**(C-D)** Sperm competition assay. Defense score represents the ability of the interested males in siring progeny when they mate with females that subsequently remated. Offense score is the reverse (see text). Curiously, in all cases where the control and KO males are significantly different (Mann-Whitney *U* test, *, *P*\*0.05), KO males have a higher fitness. **(E-F)** Linear regression of the median value from the sperm competition assay on the median sperm length. Angular transformation is applied to the defense and offense scores. Sperm length indeed has an impact of sperm competitiveness and the trend is stronger for defense than for offense.

#### Males’ ability in repressing female remating

An important attribute of male reproduction is males’ ability to repress female remating after they have successfully inseminated the female. Previous studies (Manning 1962; Chapman et al. 2003; Ram and Wolfner 2007) have revealed that the presence of sperm (known as “sperm” effect) and seminal fluid proteins can indeed elicit female post-mating responses that include egg laying and reduced receptivity to mating. Wild type females are mated to either wild type males or males from the miRNA knockout lines (Supplemental Fig.S5A). After 2.5 days, these mated females are presented with reference males and their remating rates are recorded as presented in Fig. 5A. The observations show that *mir-973*, *mir-977* and *mir-983* knockout males have an advantage over wild type males in repressing female remating.

#### Meiotic drive and male viability

Since these miRNAs are X-linked, it seems plausible that they exert the fitness effect on the X Chromosome at the expense of the Y. In other words, we ask whether there is sex-linked meiotic drive associated with these new miRNAs (Wu 1983a; Wu 1983b; Jaenike 2001). The assay of male fertility would have missed the action of meiotic drive, which affects the ratio but not the total progeny count. To score the meiotic drive, we set up the crosses: M/FM7c females × M/Y males where M represents the X Chromosome bearing a miRNA deletion (FM7c is the standard X balancer; see Methods). For control, M is the wildtype X Chromosome (designated M+).

To test for meiotic drive, we compare the abundance of FM7c/M and FM7c/Y (female and male progeny with FM7c, respectively) as the measure of distortion in sex chromosome transmission. These two genotypes would minimize the confounding fitness effect of the homozygous M/M, or the hemizygous M/Y, genotype (Supplemental Fig.S5B). Fig. 5B shows a very slight, and statistically insignificant, sex ratio distortion in *mir-975* and *mir-977*. We conclude that these new miRNAs do not impact fitness via meiotic drive.

The crosses of M/FM7c females x M/Y males also permit the measurement of these miRs’ effects on male viability. We compare the abundance of M/Y and FM7c/Y males as shown in Fig. 5C. The three younger miRs (*mir-978*, *mir-983* and *mir-984*) all appear to affect male viability and, again, the absence of these miRs improves male viability.

#### Mating success in males

While the younger miRs appear to have some adverse effects on male viability, Fig. 5D reveals the absence of effects on male mating success among all these miRNAs.

**Figure 5.**
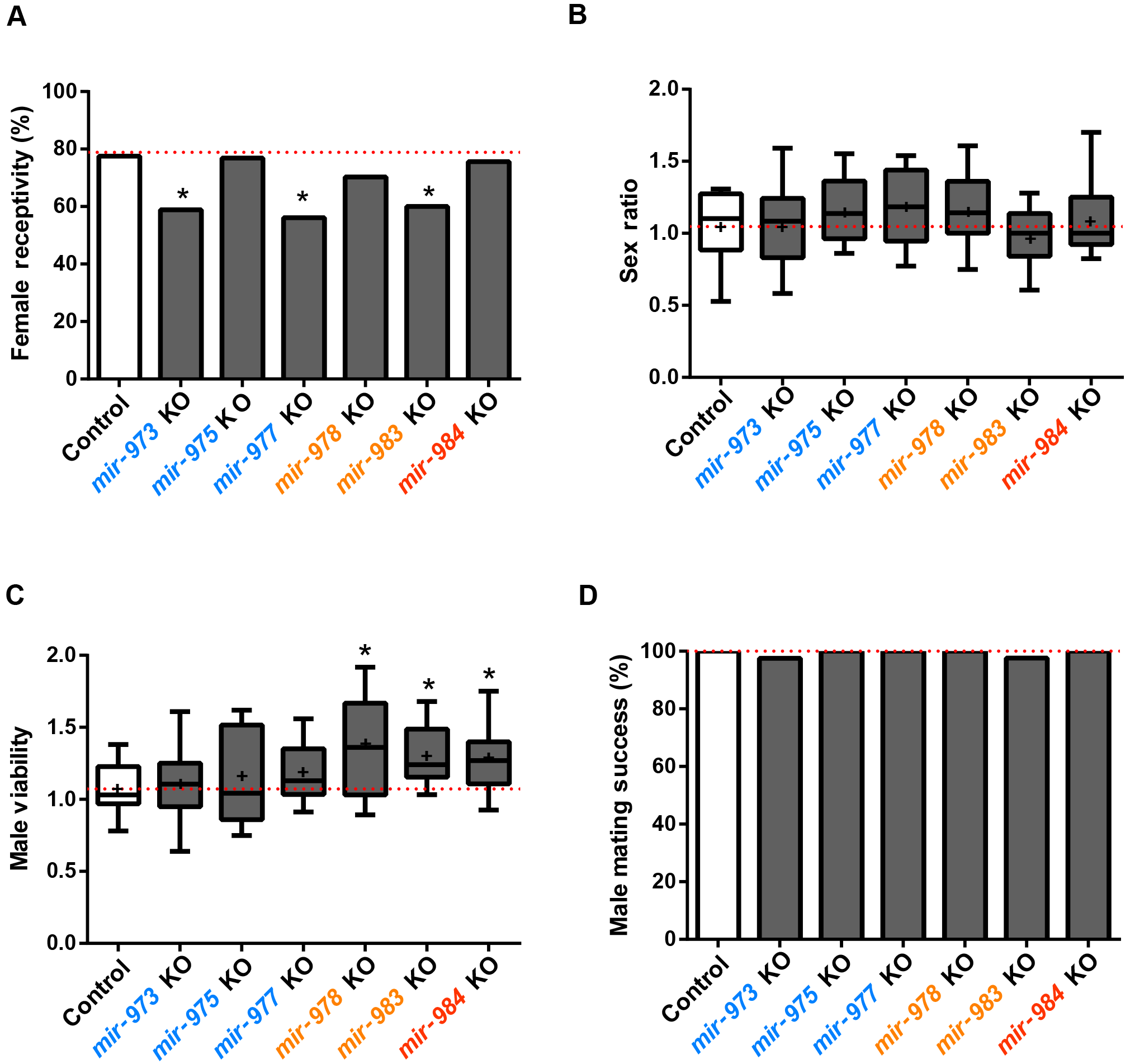
Effects on other male fitness components. **(A)** Female receptivity after mating with wild type or miRNA KO males as a measure of males’ ability to repress female remating. Lower bars here indicate higher fitness of males. In three lines, there is a significant decrease in female remating after females are mated to KO males (Chi-square test: *, *P*<0.05). **(B)** Distortion in sex chromosome transmission, a measure of miRNAs’ ability to gain a transmission advantage through the transmission of X. No difference shown is significant (Student’s *t*-test, overall *P*>0.1).**(C)** Male viability. Compared to the control, three miRNA-KO (*mir-978*, *mir-983*, and *mir-984*) males are significantly more viable (Student’s *t*-test: *, *P*<0.05). **(D)** Male mating success. Differences between miRNA mutants and controls are very small and statistically insignificant (Chi-square test, overall *P*>0.1).

### Assay of total fitness in long-term laboratory populations

The measurements of the many fitness components are summarized in Table 1. None of these fitness components reveals strong selection against miRNA deletions. In fact, males from the knockout lines of younger miRNAs tend to show higher fitness than the wild type males, thus raising the issue of these miRNAs’ existence. A relevant question is therefore whether these *de novo* miRNAs are in the process of dying. In the literature (Lyu et al. 2014), the age structure of new miRNAs suggests that younger miRNAs die frequently before reaching an old age.

**Table 1.** Relative fitness of miRNA knockout males in different components

|  | Measurements<br>in wild type males | Relative fitness of miRNA knock out males |  |  |  |  |  |  |
| --- | --- | --- | --- | --- | --- | --- | --- | --- |
|  |  | Wild type<br>(Control) | *mir-973* | *mir-975* | *mir-977* | *mir-978* | *mir-983* | *mir-984* |
| Male fertility |  |  |  |  |  |  |  |  |
| Days 1-2 | 29.3±3.3 | 1 | 1.265 | 1.165 | 1.286 | 0.935 | 0.934 | 1.111 |
| Days 3 to7 | 32.6±3.3 | 1 | 0.858 | 0.77 | 0.577* | 1.22 | 0.903 | 0.795 |
| Total | 61.9±5.8 | 1 | 1.051 | 0.957 | 0.899* | 1.075 | 0.918 | 0.943 |
| Sperm length(millimeter) | 1.883±14 | 1 | 1.008* | 1.010* | 0.973* | 1.028*** | 1.006* | 1.015** |
| Sperm competition |  |  |  |  |  |  |  |  |
| Offense | 0.958± 0.02 | 1 | 1.034 * | 0.972 | 0.963 | 1.023 * | 1.034* | 1.028 |
| Defense | 0.394± 0.05 | 1 | 1.287 | 1.341 | 0.774 | 1.298* | 1.296 | 1.049 |
| Suppression of female remating | 22.5% | 1 | 1.24* | 1 | 1.28* | 1.09 | 1.23* | 1.02 |
| Meiotic drive | 1.05±0.06 | 1 | 1.00 | 1.09 | 1.13 | 1.09 | 0.927 | 1.04 |
| Male viability | 1.07±0.04 | 1 | 1.03 | 1.07 | 1.10 | 1.29* | 1.21* | 1.20* |
| Male mating success | 100% | 1 | 0.975 | 1 | 1 | 1 | 0.976 | 1 |

**Note:** Relative fitness estimates are presented in materials and methods. (\*)*P* < 0.05, (\*\*) *P* < 0.01,(\*\*\*)*P* < 0.001.

Although the individual fitness components suggest impending gene death, it is nevertheless possible that there are other aspects of fitness that may offset the components we measured. It would be more convincing if all fitness components are measured in one single experiment. We therefore carry out the total fitness assay for two of the three younger miRNAs (< 30 Myrs in age, *mir-978* and *mir-984*). The third one (*mir-983*) has several novel properties that are analyzed in the companion study(Zhao et al. 2018). For the total fitness, the miRNA knockout and wild-type allele are assayed in laboratory populations, following the established protocols (Fig. 6A; see Methods) (Wu et al. 1989; Alipaz et al. 2005; Fang et al. 2012). In long-term populations, components of selection in aggregate are manifested as gene frequency changes (Supplemental Fig. S6). Each population with a population size of ~1800 lasts 40 generations.

Although many individual selection components favor the elimination of the *de novo* miRNAs (Table 1), the trend in gene frequency stays constant in the long-term assays (Overall *P* > 0.1, One-sample *t*-test, Fig.6B-6C). Using the maximum likelihood estimation, we obtain the selective coefficient (s) is 0.0025 and −0.005, respectively, for the *mir-978* and *mir-984* mutant. Neither is significantly different from 0 (Fig.6 D-6E, *P* > 0.05, Chi-square test). In contrast, if one uses the fitness component estimates shown in Table 1, both mutant alleles are expected to be higher than 90% after 40 generations. The apparent discrepancy between the results of selection components and total fitness is highly instructive as discussed below.

**Figure 6.**
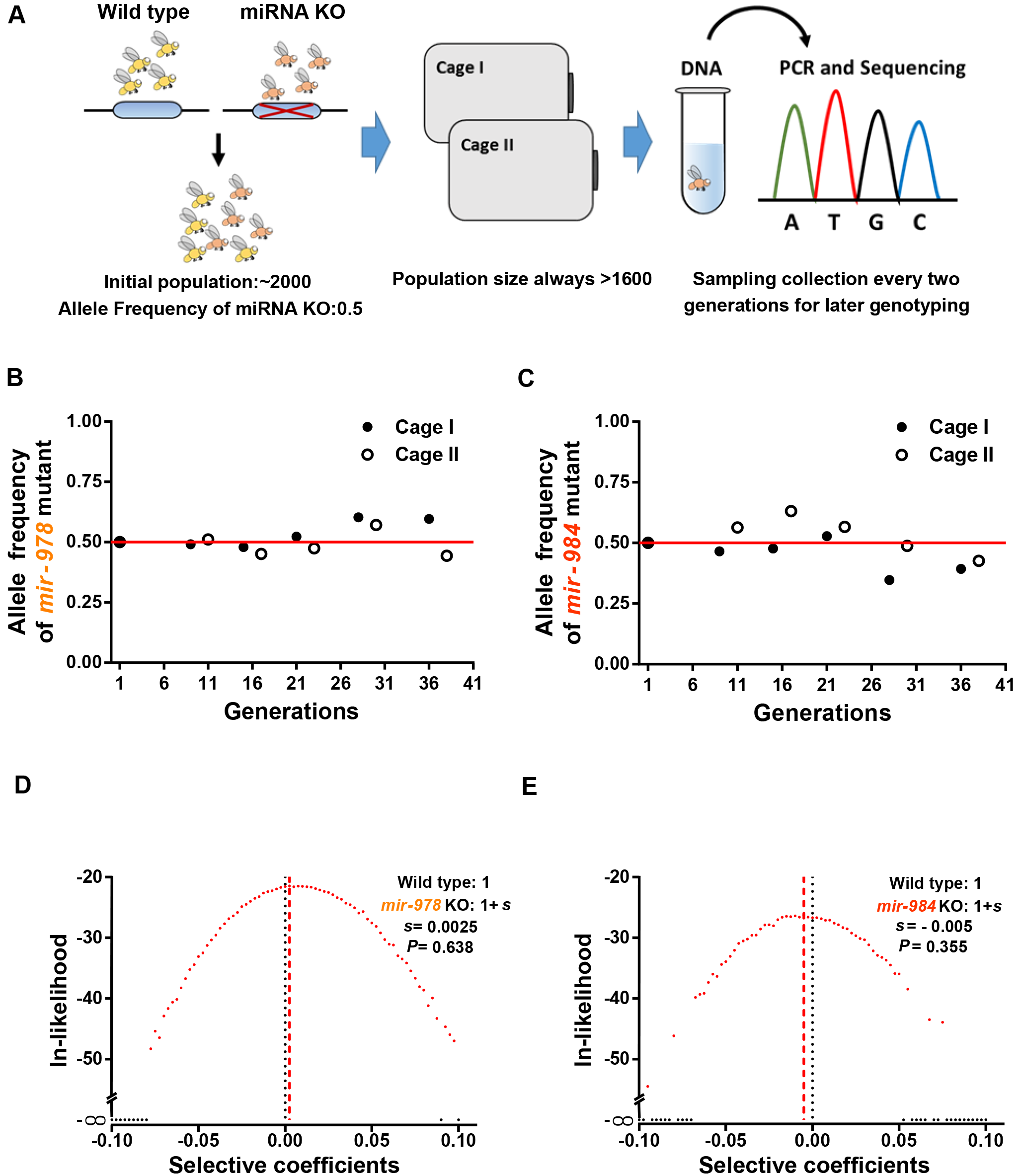
Assay for total fitness of miRNA deletions in laboratory population. **(A)** Set-up of laboratory populations. Throughout the experiment, the population size never drop below 1600. A sample of about ~ 200 flies is collected from each cage every 2 generations. Genotype identification by single fly PCR and sequencing is done on samples of various time points. **(B-C)** Allele frequency changes of miRNA KO mutants in laboratory populations over ~ 40 generations. Allele frequencies do not appear to deviate from the initial frequency of 0.5 (see D-E for details). **(D-E)** Likelihood distributions of the selective coefficient of miRNA KO mutants. The fitness of miRNA KO alleles is 1+s whereby s is expected to be negative. The maximal likelihood estimate of s is 0.0025 and −0.0005, respectively, for *mir-978* and *mir-984*. Neither is significantly different from 0 (*P*=0.638 and 0.355 by the Chi-square test).

## Discussion

The study of total fitness in long-term laboratory populations reveals little net selection suggesting that many components of fitness are not captured in Table 1. One would then ask how a single miRNA could play a role in such a diverse array of components and, intriguingly, often in opposite directions. The subject has been addressed recently by Liufu et al. who show that a typical miRNA often controls multiple phenotypes and sub-phenotypes redundantly and incoherently via many different target genes(Liufu et al. 2017). The reasons for such incoherent regulatory mechanisms are further explained in two recent publications (Chen Y 2017; Zhao et al. 2017) in terms of network stability. It is therefore plausible that new miRNAs would also impact many selection components weakly, redundantly and incoherently.

Since new genes, both coding and noncoding, are constantly born (Bai et al. 2007; Wolf et al. 2009; Berezikov et al. 2010) while the total number of genes remain steady among taxa, death soon after birth is expected. The age structure of *de novo* miRNAs, with far fewer older miRNAs than younger ones, is an indication of death (Lu et al. 2008; Lyu et al. 2014).

How and why do new genes die? For example, how does natural selection reverse its course when a previously adaptive gene becomes fitness-neutral? The newly evolved state may be more appropriately referred to as “quasi-neutrality”. While neutrality is usually equated with the loss of functionality (i.e., non-functionalization; (Li et al. 1981), genes of quasi-neutrality remain functional. Such a gene is quasi-neutral because selection for some components (e.g. male fertility) and against others (e.g. sperm competition) results in nearly zero net gain/loss in fitness. The evolution of *de novo* miRNA genes thus broadens the definition of neutral evolution. When a gene loses its selective advantages and becomes neutral or quasi-neutral, the time it takes for the gene to die would be roughly 1/u + 4Ne generations, where u is the rate of gene non-functionalization(Crow and Kimura 1970). In the case of quasi-neutrality, the selective pressure is present but averages to be zero through time and space. Theoretical study shows that the evolutionary dynamics would be quite complicated (Cvijovic et al. 2015). In particular, the duration of dying could be substantially shorter or longer than the true neutral genes. This broader range of dynamics makes it more likely to observe the process of gene dying.

This study of *de novo* genes has general implications. It shows that the evolution of a new function, in the form of either a gene or a phenotype, may often be “transiently adaptive”. Environmental pressures, either biotic or abiotic, may not stay constant and the responses should not be expected to be permanent either. This ever-shifting pressure to adapt is reminiscent of the metaphor of the “Red Queen” effect (Van Valen 1973; Salathé et al. 2008; Liow et al. 2011). In the Red Queen landscape, adaptive evolution does not always resemble the large-scale and stable adaptive changes; for example, when terrestrial woody plants invade the intertidal habitats (Xu et al. 2017a; Xu et al. 2017b) or mammals migrate to the high-altitude terrains (Yi et al. 2010). Instead, adaptations may often be transient, small-scale and local.

The Red Queen hypothesis has been extensively applied to phenotypic evolution (Dercole et al. 2006; Decaestecker et al. 2007; Morran et al. 2011). Its application to molecular evolution is more difficult because of the abundance of non-adaptive changes at the molecular level. In this study, we show that a manifestation of the Red Queen effect in molecular evolution may be the loss of previously adaptive new genes.

## Methods

### Fly stocks and miRNA mutant generation

miRNA mutants are generated using the TALEN-based gene KO system (Boch 2011). *mir-973*, *mir-975*,*mir-978* and *mir-984* knockout lines are made in this study and *mir-977*, *mir-983* knockout lines are made earlier by us(Lu et al. 2018; Zhao et al. 2018). TALEN binding sites are designed by TAL Nucleotide Targeter 2.0 (http://tale-nt.cac.cornell.edu). The TALEN plasmids are constructed by ViewSolid Biotech. Plasmids are transcribed *in vitro* to mRNAs using mMESSAGE mMACHINE T7 Ultra. A detailed cross scheme and related fly stocks are shown in Supplemental Fig.S1. TALEN binding sites are listed in Supplemental Table S2. All flies are raised at 25°C on a standard sugar-yeast-agar medium under a 12:12 hr light/dark cycle.

miRNA mutants are detected by PCR and verified by qRT-PCR (Fig.1C to H). Three batches of 50 testes are used independently as biological replicates for qRT-PCR assays. For each replicate, total RNA is extracted using the Ambion TRIzol^®^ Reagent (code No. 15596018). qRT-PCR of miRNAs is conducted using stem-loop reverse transcription (Chen et al. 2005) followed by TaqMan PCR analysis using the miRNA UPL (Roche Diagnostics) probe assay protocol (Varkonyigasic et al. 2007). For *mir-978* expression analysis, TaqMan probes are used to get higher specificity for detecting its two iso-miRNAs. Relative expression levels of miRNAs are obtained by the 2^−ΔΔCT^method (Livak and Schmittgen 2001). 2S RNA (for *mir-973*, *mir-975*, *mir-977*, *mir-983*, and *mir-984*) and u14 (for miR-978-3p.1 and miR-978-3p.2) are used as endogenous controls (Relevant primers are listed in Supplemental Table S3).

### miRNA expression estimates

Small RNA-seq data are obtained from the NCBI Gene Expression Omnibus (GEO, https://www.ncbi.nlm.nih.gov/geo/), described in previous studies(Czech et al. 2008; Rozhkov et al. 2010; Czech et al. 2012). Sequences of precursors and mature miRNAs of *D. melanogaster* are obtained from miRBase(Kozomara and Griffiths-Jones 2014), mature miRNAs expression levels(Supplemental Table S1) are calculated using all reads mappable to the mature sequences and scaling as Reads Per Million (RPM),using miRDeep2 (version 2.0.0.7)(Friedländer et al. 2012).

### RNA-seq analyses

#### Sample preparation and differential gene expression estimates

For *mir-978* and *mir-984* KO males, two batches of testes are dissected independently as biological replicates for each pair of wild type and miRNA deletion mutants. Total RNA is extracted from 30-50 pairs of testes from three- to five-day-old males using TRIzol. RNA libraries are prepared and sequenced on Illumina HiSeq 2000 at BGI (http://www.genomics.cn/index). Reference *Drosophila* genome is download from FlyBase (*D. melanogaster* genome r6.04). Data were deposited in Genome Sequence Archive (http://gsa.big.ac.cn/) under accession CRA000857. For *mir-983* KO males, sequencing data are retrieved from GEO under accession GSE110086(Zhao et al. 2018). Gene expression levels are calculated using Reads Per Kilobase per Million mapped reads (RPKM), using TopHat (Trapnell et al. 2009) and SeqMonk; genes with log2 (RPKM) <1 are filtered out as sequencing error. Differentially expressed genes between two samples with replicates are identified using DESeq2 with *P* < 0.05 (Love et al. 2014). To survey the Impact of miRNAs on target gene expression, nontargets set in Fig 2 is the gene pool containing all expressed genes without focused miRNA targets. Confirmatory analyses with refining background control (with similar expression and 3’UTR length) and additional control (target pool of a neighboring *de novo* miRNA not affected by the deletions) are shown in Supplemental Fig S7 and Supplemental Fig S8, respectively.

#### miRNA target prediction

3’UTR sequences are retrieved from FlyBase (Attrill et al. 2016). Longest UTR of each gene is chosen for target prediction. Target genes of miR-975-5p, miR-983-5p, miR-984-5p and miR-978-3p are predicted using the TargetScan algorithm with default settings (Ruby et al. 2007). All predicted targets with 8mer, 7mer-m8 and 7mer-1A sites are used to examine the repression effect. Since these miRNA are recently evolved, target site conservation would not be a useful criterion. Here we use ‘seed matches’ for target prediction without conservation considerations.

### Male fertility assays

To survey male fertility (Fig.3 and Supplemental Fig.S3), we count the number of eggs (denoted x), and progeny (1^st^ instar larvae, denoted y) produced by wild type females mate to experimental (miRNA mutant or control) males for 7 days (Supplemental Fig.S2). Male fertility in days 1-2 is y_1_+y_2_; male fertility in days 3-7 is y_3_+y_4_+y_7_. We score ~12 single pair replicates per assay. Each female in a pair is exposed to a male for 2 hours.

### Sperm assays

#### Sperm length

We measured *Drosophila* sperm length as described in the Fiona M. Hunter and T.R. Birkhead (Hunter and Birkhead 2002). Briefly, we remove seminal vesicles from five-to six-day-old males and place them in drops of phosphate-buffered saline (PBS) on glass slides. Sperm is liberated from tissues by poking with a needle. We measure sperm length using supporting software LAS V3.8 on a Leica DMI4000 microscope. We measure 9 to11 individuals per genotype and 9 to 13 sperm cells per individual.

#### Sperm competition

Sperm competition assays are conducted as described in the Shu-Dan Yeh *et al* (Yeh et al. 2013). Briefly, double-mating experiments (Supplemental Fig.S4) are set up for each miRNA mutant and control line. Virgin *w*^*1118*^ females are mated to reference males for 2h and then mated to experimental males (miRNA KO or control) for 12h after 56h to measure offensive ability. Conversely, *w*^*1118*^ females are mated to experimental males for 2h and then to the reference males for 12h after 56h for the defense assay. Each assay had 40 replicates. Eye color of progeny is used as a marker for evaluations of mating success and paternity identification. Females who have both red-eyed and white-eyed progeny have successfully mated with both experimental and reference male. The red-eyed progeny (y) are from reference males, while white-eye progeny (x) are from experimental fathers. Sperm competitive ability is estimated using the score P = x /(x + y). Angular transformation is applied to the defense and offense score for linear regression.

Compared to the original protocol, the only change in our assay is that we use *w*^*1118*^/Y; *miniwhite-UASeGFP*/*miniwhite-UASeGFP*, a red-eyed strain with the insertion of *mini white* at the 51D position on chromosome 2 via the PhiC31 site-specific chromosomal integration system.

### Mating success and males’ ability to repress female re-mating

To survey mating success and males’ ability to repress female re-mating, wild type females are mated to either wild type or miRNA knockout males. After 2.5 days, these mated females are presented with reference males (Supplemental Fig. S5A). Mating rates of F0 females are inferred from progeny types. We classify F0 female mating success, number of females (x) with white-eye progeny only (successful fertilization by experimental male only), number of female (y) with both white-eyed and red-eyed progeny (successful fertilization by experimental and reference males, Supplemental Fig.S5A). Male mating success is then (x + y)/N_replicates_, while female receptivity is y/(x + y).

### Male viability and meiotic drive

We cross balanced mutant or control females (FM7c/M) to hemizygous mutant or control males (M/Y, Supplemental Fig. S5B). In the F1 progeny for each cross, the ratio of M/Y to FM7c/Y males reflects male viability of wild type or miRNA knockouts compared to balancer males. The ratio of M/M to FM7c/M females reflects female viability of wild type or miRNA knockouts compared to balancer females, which can be used as an additional control to rule out the possibility of off-target effects (Supplemental Table S4; Supplemental text S2). Although the balancers can affect viability, this indirect comparison controls for the balancer influence. The abundance of FM7c/M *vs* FM7c/Y (female and male progeny with FM7c) reflects distortion in sex chromosome transmission.

### Estimates of relative fitness

To estimate relative fitness statistics shown in Table1, we denote wild type values as w and those of deletion males as m. We set the fitness of non-mutant genotypes as 1. Fitness of miRNA deficient males is then m/w for most measures. The exception is female re-mating suppression. In this case, the suppression intensity is 1-w or 1-m for wild type and mutant respectively. To reflect the relative fitness in suppression of female re-mating, we use (w - m)/w + 1, where (w - m)/w is the relative fitness gain of miRNA knockout over wild type.

### Allele frequency estimation in population cage assays

Population cage assays are performed as described in a previous study of ours (Wu et al. 1989). Each competition assay includes two replicates. Each cage is started with 1000 miRNA deletion and 1000 wild type flies with the sex ratio 1:1. The populations are maintained at 25°C with overlapping generations. Population size is maintained at about 1600~2000 flies for every generation. Sampling is done every two generations by collecting flies that emerged from several food bottles. At specific generations, 96 males and 96 females are sampled randomly to extract DNA and perform single fly PCR. From the PCR results, we define each fly’s genotype (Supplemental Fig. S6) and estimate the miRNA mutant frequency for that generation.

### Maximum likelihood estimates of selection coefficients

The selection coefficient estimates are done according to a previous study of ours (Fang et al. 2012).

#### Simulation of populations under the Wright-Fisher model

Each simulation involves 10,000 iterations with fixed selection coefficient (s) values. Each iteration is ran for 40 generations with population sizes fixed at 800 males and 800 females each generation. Each generation is divided into selection (male only) and reproduction, fitness is 1 for wild type and 1+s for deletion males. Female fitness is 1 for both genotypes. During random mating, male and female haplotypes are randomly selected from the population to make 800 males and 800 females for the next generation.

#### Sampling and likelihood estimates

Simulation of sampling involves 10,000 iterations. Sampling is done in the generations as shown in Fig. 6B and C, each sample contains 90 males and 90 females; sampling time is after selection stage and before reproduction stage. Allele frequencies of the miRNA mutant are recorded from these sampled flies. These recorded allele frequencies are assign to 40 bins (0-.025, .025-.05,…, .975-1) to create distributions for the sampling generations. For each fixed *s*, likelihood ratio is ln (No. of bins containing observed frequencies/ No. of iterations).

Simulations are ran for 80 sets of s values (−0.1~0.1), and the maximum likelihood estimate for each miRNA deletion’s selection coefficient is the *s* value that yielded the maximum likelihood ratio.

## Data access

RNA-seq data for wild type, *mir-978KO* and *mir-984* KO testis have been submitted to the BIG Data Center in Beijing Institute of Genomics (http://bigd.big.ac.cn/) with accession number CRA000857. RNA-seq data for wild type and *mir-983* KO testis are retrieved from GEO under accession GSE110086.

## Acknowledgments

We thank all members in Wu lab for helpful comments and ideas sharing.

We thank Huang Yumei, Ma Fuqiang, Shang Rui and Shen Xu, Sun Yat-sen University, for generous help with data collection of population cages; Li Xiang, Sun Yat-sen University, for help in male fertility and sperm competition assays; Shang Rui for help in sperm length data re-analysis. We thank Lu Jian, Peking University, Shi Shuhua, Sun Yat-sen University,for comments on previous versions of the manuscript.

This work is supported by the National Science Foundation of China (31730046, 91231117, 31130069), the 985 Project (33000-18821105), the National Basic Research Program (973 Program) of China (2014CB542006).

## Author contributions

Conceived and designed the experiments: Guang-An Lu, Zhongqi Liufu, Tian Tang, Chung-I Wu. Performed the experiments: Guang-An Lu, Yixin Zhao (Mutant generation for *mir-984*), Zhongqi Liufu (Mutant generation for *mir-975*). Analyzed data: Guang-An Lu, Yixin Zhao (Transcriptome analysis), Ao Lan (Likelihood estimates for the population cage results). Wrote the paper: Guang-An Lu, Jin Xu, Haijun Wen and Chung-I Wu.

## References

Alipaz JA, Fang S, Osada N, Wu C. 2005. Evolution of Sexual Isolation during Secondary Contact: Genotypic versus Phenotypic Changes in Laboratory Populations. The American Naturalist 165: 420–428.

Attrill H, Falls K, Goodman JL, Millburn GH, Antonazzo G, Rey AJ, Marygold SJ, FlyBase c. 2016. FlyBase: establishing a Gene Group resource for Drosophila melanogaster. Nucleic Acids Res 44: D786–792.

Bai Y, Casola C, Feschotte C, Betran E. 2007. Comparative genomics reveals a constant rate of origination and convergent acquisition of functional retrogenes in Drosophila. Genome Biology 8: 1–9.

Bartel DP. 2004. MicroRNAs: genomics, biogenesis, mechanism, and function. Cell 116: 281–297.

Berezikov E, Liu N, Flynt AS, Hodges E, Rooks M, Hannon GJ, Lai EC. 2010. Evolutionary flux of canonical microRNAs and mirtrons in Drosophila. Nature Genetics 42: 6–10.

Boch J. 2011. TALEs of genome targeting. Nature Biotechnology 29: 135–136.

Cassidy JJ, Jha AR, Posadas DM, Giri R, Venken KJT, Ji J, Jiang H, Bellen HJ, White KP, Carthew RW. 2013. miR-9a Minimizes the Phenotypic Impact of Genomic Diversity by Buffering a Transcription Factor. Cell 155: 1556–1567.

Chapman T, Bangham J, Vinti G, Seifried B, Lung O, Wolfner MF, Smith HK, Partridge L. 2003. The sex peptide of Drosophila melanogaster: female post-mating responses analyzed by using RNA interference. Proc Natl Acad Sci U S A 100: 9923.

Chen C, Ridzon D, Broomer A, Zhou Z, Lee DH, Nguyen JT, Barbisin M, Xu NL, Mahuvakar VR, Andersen MR. 2005. Real-time quantification of microRNAs by stem-loop RT-PCR. Nucleic Acids Research 33: e179.

Chen S, Krinsky BH, Long M. 2013. New genes as drivers of phenotypic evolution Nature Reviews Genetics 14: 745–745.

Chen Y AS, Shen Y, Wu C-I. 2017. From foodwebs to gene regulatory networks (GRNs): weak repressions by microRNAs confer system stability. bioRxiv: doi:10.1101/176701.

Crow JF, Kimura M. 1970. An Introduction to Population Genetics Theory. Burgess Minneapolis.

Cvijovic I, Good BH, Jerison ER, Desai MM. 2015. Fate of a mutation in a fluctuating environment. Proceedings of the National Academy of Sciences of the United States of America 112: E5021–E5028.

Czech B, D’Alterio C, Jones DL, Levine E, Toledano H. 2012. The let-7-Imp axis regulates ageing of the Drosophila testis stem-cell niche. Nature 485: 605–610.

Czech B, Malone CD, Zhou R, Stark A, Schlingeheyde C, Dus M, Perrimon N, Kellis M, Wohlschlegel JA, Sachidanandam R. 2008. An endogenous small interfering RNA pathway in Drosophila. Nature 453: 798–802.

Decaestecker E, Gaba S, Raeymaekers JA, Stoks R, Van KL, Ebert D, De ML. 2007. Host-parasite ‘Red Queen’ dynamics archived in pond sediment. Nature 450: 870.

Dercole F, Ferrière R, Gragnani A, Rinaldi S. 2006. Coevolution of slow-fast populations: evolutionary sliding, evolutionary pseudo-equilibria and complex Red Queen dynamics. Proceedings Biological Sciences 273: 983–990.

Domazetlošo T, Carvunis AR, Albà MM, Šestak MS, Bakarić R, Neme R, Tautz D. 2017. No evidence for phylostratigraphic bias impacting inferences on patterns of gene emergence and evolution. Molecular Biology and Evolution 34: 843–856.

Fang S, Yukilevich R, Chen Y, Turissini DA, Zeng K, Boussy IA, Wu CI. 2012. Incompatibility and competitive exclusion of genomic segments between sibling Drosophila species. PLoS Genet 8: e1002795.

Friedländer MR, Mackowiak SD, Li N, Chen W, Rajewsky N. 2012. miRDeep2 accurately identifies known and hundreds of novel microRNA genes in seven animal clades. Nucleic Acids Research 40: 37–52.

Gnad F, Parsch J. 2006. Sebida: a database for the functional and evolutionary analysis of genes with sex-biased expression. Bioinformatics 22: 2577–2579.

Griffiths RC, Mckechnie SW, Mckenzie JA. 1982. Multiple mating and sperm displacement in a natural population of Drosophila melanogaster. Tagtheoretical & Applied Geneticstheoretische Und Angewandte Genetik 62: 89–96.

Gubala AM, Schmitz JF, Kearns MJ, Vinh TT, Bornberg-Bauer E, Wolfner MF, Findlay GD. 2017. The goddard and saturn genes are essential for Drosophila male fertility and may have arisen de novo. Molecular Biology & Evolution 34: 1066–1082.

Haldane JBS. 1932. The Causes of Evolution Harper and Bros, New York

Hunter FM, Birkhead TR. 2002. Sperm Viability and Sperm Competition in Insects. Current Biology 12: 121–123.

Jacob F. 1977. Evolution and tinkering. Science 196: 1161–1166.

Jaenike J. 2001. SEX CHROMOSOME MEIOTIC DRIVE. Annual Review of Ecology, Evolution, and Systematics 32: 25–49.

Jones B, Clark AG. 2003. Bayesian Sperm Competition Estimates. Genetics 163: 1193–1199.

Kozomara A, Griffiths-Jones S. 2014. miRBase: annotating high confidence microRNAs using deep sequencing data. Nucleic Acids Research 42: D68.

Lüpold S, Manier MK, Puniamoorthy N, Schoff C, Starmer WT, Luepold SHB, Belote JM, Pitnick S. 2016. How sexual selection can drive the evolution of costly sperm ornamentation. Nature 533: 535–538.

Li W-H. 1982. Evolutionary change of duplicate genes. Isozomes: Current Topics in Medical and Biological Research VI: 55–92.

Li WH, Gojobori T, Nei M. 1981. Pseudogenes as a paradigm of neutral evolution. Nature 292: 237–239.

Liow LH, Van VL, Stenseth NC. 2011. Red Queen: from populations to taxa and communities. Trends in Ecology & Evolution 26: 349–358.

Liufu Z, Zhao Y, Guo L, Miao G, Xiao J, Lyu Y, Chen Y, Shi S, Tang T, Wu CI. 2017. Redundant and incoherent regulations of multiple phenotypes suggest microRNAs’ role in stability control. Genome Research 27: 1665–1673.

Livak KJ, Schmittgen TD. 2001. Analysis of relative gene expression data using real-time quantitative PCR and the 2(-Delta Delta C(T)) Method. Methods 25: 402–408.

Love MI, Huber W, Anders S. 2014. Moderated estimation of fold change and dispersion for RNA-seq data with DESeq2. Genome biology 15: 550.

Lu GA, Zhao Y, Liufu Z, Wu CI. 2018. On the possibility of death of new genes-evidence from the deletion of de novo microRNAs. BMC Genomics 19: 388.

Lu J, Shen Y, Carthew RW, Wang SM, Wu C. 2010. Reply to evolutionary flux of canonical microRNAs and mirtrons inDrosophila. Nature Genetics 42: 9–10.

Lu J, Shen Y, Wu Q, Kumar S, He B, Shi S, Carthew RW, Wang SM, Wu C. 2008. The birth and death of microRNA genes in Drosophila. Nature Genetics 40: 351.

Lynch M, Conery JS. 2003. The origins of genome complexity. Science 302: 1401–1404.

Lyu Y, Shen Y, Li H, Chen Y, Guo L, Zhao Y, Hungate E, Shi S, Wu C-I, Tang T. 2014. New MicroRNAs in Drosophila—Birth, Death and Cycles of Adaptive Evolution. Plos Genetics 10: 229–231.

Manning A. 1962. A Sperm Factor Affecting the Receptivity of Drosophila Melanogaster Females. Nature 194: 252–253.

Mclysaght A, Hurst LD. 2016. Open questions in the study of de novo genes: what, how and why. Nature Reviews Genetics 17: 567.

Meunier J, Lemoine F, Soumillon M, Liechti A, Weier M, Guschanski K, Hu H, Khaitovich P, Kaessmann H. 2013. Birth and expression evolution of mammalian microRNA genes. Genome Research 23: 34–45.

Miller GT, Pitnick S. 2002. Sperm-female coevolution in Drosophila. Science 298: 1230–1233.

Mohammed J, Bortolamiolbecet D, Flynt AS, Gronau I, Siepel A, Lai EC. 2014. Adaptive evolution of testis-specific, recently evolved, clustered miRNAs in Drosophila. RNA 20: 1195–1209.

Morran LT, Schmidt OG, Gelarden IA, Nd PR, Lively CM. 2011. Running with the Red Queen: host-parasite coevolution selects for biparental sex. Science 333: 216–218.

Moyers BA, Zhang J. 2015. Phylostratigraphic bias creates spurious patterns of genome evolution. Molecular Biology & Evolution 32: 258.

Moyers BA, Zhang J. 2016. Evaluating Phylostratigraphic Evidence for Widespread De Novo Gene Birth in Genome Evolution. Molecular Biology & Evolution 33: 1245.

Moyers BA, Zhang J. 2017. Further simulations and analyses demonstrate open problems of phylostratigraphy Genome Biol Evol 9: 1519–1527.

Ohno S. 1970. Evolution by Gene Duplication. Springer Berlin Heidelberg.

Parker GA. 1970. SPERM COMPETITION AND ITS EVOLUTIONARY CONSEQUENCES IN THE INSECTS. Biological Reviews 45: 525–567.

Pattarini JM, Starmer WT, Bjork A, Pitnick S. 2006. MECHANISMS UNDERLYING THE SPERM QUALITY ADVANTAGE IN DROSOPHILA MELANOGASTER. Evolution 60: 2064–2080.

Petrov DA, Hartl DL. 1998. HIGH RATE OF DNA LOSS IN THE DROSOPHILA MELANOGASTER AND DROSOPHILA VIRILIS SPECIES GROUPS. Molecular Biology and Evolution 15: 293–302.

Petrov DA, Lozovskaya ER, Hartl DL. 1996. High intrinsic rate of DNA loss in Drosophila. Nature 384: 346–349.

Posadas DM, Carthew RW. 2014. MicroRNAs and their roles in developmental canalization. Current Opinion in Genetics & Development 27: 1–6.

Ram KR, Wolfner MF. 2007. Sustained Post-Mating Response in Drosophila melanogaster Requires Multiple Seminal Fluid Proteins. PLOS Genetics 3: e238.

Rozhkov NV, Aravin AA, Zelentsova ES, Schostak NG, Sachidanandam R, Mccombie WR, Hannon GJ, Evgen’Ev MB. 2010. Small RNA-based silencing strategies for transposons in the process of invading Drosophila species. Rna-a Publication of the Rna Society 16: 1634.

Ruby JG, Stark A, Johnston WK, Kellis M, Bartel DP, Lai EC. 2007. Evolution, biogenesis, expression, and target predictions of a substantially expanded set of Drosophila microRNAs. Genome Research 17: 1850–1864.

Salathé M, Kouyos RD, Bonhoeffer S. 2008. The state of affairs in the kingdom of the Red Queen. Trends in Ecology & Evolution 23: 439.

Schlötterer C. 2015. Genes from scratch – the evolutionary fate of de novo genes. Trends in Genetics 31: 215–219.

Tautz D, Domazetlošo T. 2011. The evolutionary origin of orphan genes. Nature Reviews Genetics 12: 692–702.

Trapnell C, Pachter L, Salzberg SL. 2009. TopHat: discovering splice junctions with RNA-Seq. Bioinformatics 25: 1105–1111.

Van Valen L. 1973. A new evolution law. Evolutionary Theory 1: 1–30.

Varkonyigasic E, Wu R, Wood M, Walton EF, Hellens RP. 2007. Protocol: a highly sensitive RT-PCR method for detection and quantification of microRNAs. Plant Methods 3: 12–12.

Wolf YI, Novichkov PS, Karev GP, Koonin EV, Lipman DJ. 2009. The universal distribution of evolutionary rates of genes and distinct characteristics of eukaryotic genes of different apparent ages. Proceedings of the National Academy of Sciences of the United States of America 106: 7273–7280.

Wu C. 1983a. VIRILITY DEFICIENCY AND THE SEX-RATIO TRAIT IN DROSOPHILA PSEUDOOBSCURA. I. SPERM DISPLACEMENT AND SEXUAL SELECTION. Genetics 105: 651–662.

Wu C. 1983b. VIRILITY DEFICIENCY AND THE SEX-RATIO TRAIT IN DROSOPHILA PSEUDOOBSCURA. II. MULTIPLE MATING AND OVERALL VIRILITY SELECTION. Genetics 105: 663–679.

Wu CI, Shen Y, Tang T. 2009. Evolution under canalization and the dual roles of microRNAs: a hypothesis. Genome Research 19: 734–743.

Wu CI, True, Amp JR, Johnson N. 1989. Fitness reduction associated with the deletion of a satellite DNA array. Nature 341: 248–251.

Wu DD, Zhang YP. 2013. Evolution and function of de novo originated genes. Molecular Phylogenetics & Evolution 67: 541.

Xu S, He Z, Guo Z, Zhang Z, Wyckoff GJ, Greenberg A, Wu CI, Shi S. 2017a. Genome-Wide Convergence during Evolution of Mangroves from Woody Plants. Molecular Biology & Evolution 34: 1008.

Xu S, He Z, Zhang Z, Guo Z, Guo W, Lyu H, Li J, Yang M, Du Z, Huang Y et al. 2017b. The origin, diversification and adaptation of a major mangrove clade (Rhizophoreae) revealed by whole-genome sequencing. National Science Review 4: 721–734.

Yeh SD, Chan C, Ranz JM. 2013. Assessing differences in sperm competitive ability in Drosophila. Journal of Visualized Experiments: e50547–e50547.

Yi X, Liang Y, Huertasanchez E, Jin X, Zha XPC, Pool JE, Xu X, Jiang H, Vinckenbosch N, Korneliussen TS. 2010. Sequencing of Fifty Human Exomes Reveals Adaptation to High Altitude. Science 329: 75.

Zhang J. 2013. Evolution by gene duplication: an update. Trends in Ecology & Evolution 18: 292–298.

Zhao L, Saelao P, Jones CD, Begun DJ. 2014. Origin and spread of de novo genes in Drosophila melanogaster populations. Science 343: 769–772.

Zhao Y, Shen X, Tang T, Wu C-I. 2017. Weak Regulation of Many Targets Is Cumulatively Powerful—An Evolutionary Perspective on microRNA Functionality. Mol Biol Evol 34(12): 3041–3046.

Zhao Y, Yang H, Lin P, Liufu Z, Lu GA, Xu J, Tang T, Wen H, Wu CI. 2018. Testing the Red Queen hypothesis on de novo new genes - Run or die in the evolution of new microRNAs. bioRxiv: doi:10.1101/345769.

